# Syrbactin-class dual constitutive- and immuno-proteasome inhibitor TIR-199 impedes myeloma-mediated bone degeneration *in vivo*

**DOI:** 10.1101/2021.10.27.466109

**Authors:** Vasudha Tandon, Ruturajsinh M Vala, Albert Chen, Robert L Sah, Hitendra M Patel, Michael C Pirrung, Sourav Banerjee

## Abstract

Proteasome-addicted neoplastic malignancies present a considerable refractory and relapsed phenotype with patients exhibiting drug-resistance and high mortality rates. To counter this global problem, novel proteasome-based therapies are being developed. In the current study we extensively characterize TIR-199, a syrbactin-class proteasome inhibitor derived from a plant virulence factor of bacterium *Pseudomonas syringae pv syringae.* We report that TIR-199 is a potent constitutive and immunoproteasome inhibitor, capable of inducing cell death in multiple myeloma, triple-negative breast cancer and non-small cell lung cancer lines, effectively inhibit proteasome in primary myeloma cells of refractory patients, and bypass the PSMB5 A49T+A50V bortezomib-resistant mutant. TIR-199 treatment leads to accumulation of canonical proteasome substrates in cells, it is specific, and does not inhibit 50 other enzymes tested in vitro. The drug exhibits synergistic cytotoxicity in combination with proteasome-activating-kinase DYRK2 inhibitor LDN192960. Furthermore, low dose TIR-199 exhibits in vivo activity in delaying myeloma-mediated bone degeneration in a mouse xenograft model. Together, our data indicates that proteasome inhibitor TIR-199 could indeed be a next generation drug within the repertoire of proteasome-based therapeutics with a potential to target relapsed and refractory proteasome-addicted neoplasia.

## Introduction

The proteolytic subunits β1/PSMB1/caspase-like, β2/PSMB2/tryptic-like, and β5/PSMB5/chymotryptic-like, represents the catalytic core of the 20S constitutive proteasome(1). The 20S catalytic core associates with 19S regulator complex to form the constitutive 26S proteasome(2). In vertebrates, specific catalytically active β subunits, i.e., β1i/LMP2/PSMB9, β2i/MECL1, and β5i/LMP7/PSMB8, are incorporated into the 20S core proteasome upon interferon IFNγ-mediated induction to form immunoproteasome(3). The constitutive 26S proteasome degrades majority of cellular proteins in eukaryotes(1) while the immunoproteasome generates peptides from antigens to present to cytotoxic T cells(4). Over the years, the proteasome has been pharmacologically targeted for the treatments of multiple myeloma(1) and mantle-cell lymphoma(5). FDA-approved proteasome inhibitor bortezomib directly inhibits the 20S core of the proteasome and have significantly improved life-expectancy of multiple myeloma patients(6), although with reported side effects and eventual drug-resistance(7). Bortezomib exhibits highest binding affinity to the PSMB5 subunit of the 20S proteasome and unfortunately, PSMB5 frequently accumulates mutations and/or increases copy numbers thereby triggering bortezomib-resistance(8,9). To circumvent the issues of bortezomib resistance and toxicity, immunoproteasome inhibitors like ONX-0914 or dual 26S and immune-proteasome inhibitors like oprozomib are being developed preclinically(10,11). ONX-0914 preferentially binds to the PSMB8 subunit of the immunoproteasome and bypasses bortezomib-resistance in myeloma cell lines(12). However, both these epoxyketone class molecules exhibit similar binding mechanism to carfilzomib which eventually might lead to resistance(13). Furthermore, various recent studies including our own have shown that multiple kinases can phosphorylate and regulate the activity of the constitutive proteasome(14–17). Interestingly, targeting one such proteasome regulating kinase DYRK2 *in vivo* dramatically reduced tumour burden in multiple myeloma and triple negative breast cancer (TNBC) mouse models(18–21). In fact DYRK2 inhibitor LDN192960 significantly reduces proteasome activity, impedes systemic bone degeneration in immunocompetent mouse myeloma models(20) and induces synergistic cytotoxicity in cancer cells in combination with proteasome inhibitors(18,20). Pharmacological targeting of such upstream kinases has indeed opened up new possibilities in proteasome-based therapeutics (18,20). Therapeutic targeting of the 26S proteasome could also be beneficial for solid tumours like triple-negative breast cancer patients (22,23) and in some cases for KRAS G12D lung cancer patients (24). On a similar note, immunoproteasome inhibitors are also being developed to target various autoimmune, cancer, and inflammatory diseases (25). Interestingly, various groups have reported that a simultaneous constitutive and immuno-proteasome inhibition could in fact have an additive/synergistic effect in alleviating refractory/relapsed multiple myeloma progression (12,26). Thus, a dual constitutive and immuno-proteasome inhibitor could be a major therapeutic blockbuster in the future.

In 2008, a plant virulence factor of bacterium *Pseudomonas* was found to directly bind and inhibit the proteasome(27). The virulence factor derivatives called syrbactins are a family of bacterial, macrocyclic, non-ribosomal peptide natural products which reacts covalently with the catalytic Thr1 residue of proteasome peptidase subunits (27–29). Upon further structure-activity-relationship studies, syrbactin analog TIR-199 was developed and found to inhibit proteasome activity(30) and induce cell death in myeloma, mantel-cell lymphoma and neuroblastoma celllines (31). Preliminary studies show that TIR-199 is active *in vivo* and arrests tumor cell growth in mice (30,31). The current work builds on the initial studies with TIR-199 and shows that TIR-199 is indeed a dual constitutive and immuno-proteasome inhibitor with high inhibitory activity against PSMB5 and PSMB8 subunits and can bypass bortezomib resistant mutant of PSMB5. TIR-199 can induce cell death in myeloma, triple-negative breast cancer and non-small cell lung cancer and can trigger synergistic cytotoxicity in combination with DYRK2 inhibitor LDN192960. Although a highly hydrophobic natural product derivative, TIR-199 is specific and does not inhibit 50 kinases tested *in vitro* and effectively impedes myeloma-mediated bone degeneration *in vivo* at low dose. Overall, we report a highly specific, bioactive syrbactin based proteasome inhibitor with potent anti-cancer properties.

## Materials and Methods

### Materials

Antibody against 20S proteasome (#BML-PW8195) and purified human 20S proteasome (#BML-PW8720) were from Enzo Lifesciences. Flag-M2 antibody (#F3165) was from MerckMillipore. GAPDH antibody (#CB1001500UG) was from Calbiochem. Antibodies against p62 (#5114), p21 (#2947), beta-tubulin (#2128), and IκBα (#9242) were from Cell Signalling.

### General methods

All recombinant DNA procedures, electrophoresis, Coomassie-staining, immunoblotting, and FLAG affinity purification were performed using standard protocols. DNA constructs used for transfection were purified from *Escherichia coli* DH5α using Macherey-Nagel NucleoBond^®^ Xtra Maxi kits according to the manufacturer’s protocol. PSMB5 A49T+A50V (A108T+A109V in full length PSMB5) mutagenesis was performed using the QuikChange^®^ site-directed mutagenesis method (Stratagene) with Q5 polymerase (New England Biolabs). All DNA constructs were verified by DNA sequencing. For qRT-PCR analysis, total RNA from HEK293 cells were isolated using the NucleoSpin RNA kit (Macherey-Nagel, Bethlehem, PA). cDNA was synthesized using the iScript kit (Bio-Rad). qRT-PCR analysis was performed using the SYBR^®^ Premix Ex Taq™ II (Takara) on Applied Biosystems 7500 Real-Time PCR System. Data were normalized to corresponding GAPDH levels. Primers used for hGAPDH (Forward: ACATCGCTCAGACACCATG; Reverse: TGTAGTTGAGGTCAATGAAGGG), hPSMB8 (Forward: CGGGTAGTGGGAACACTTATG; Reverse: TTGACAACGCCTCCAGAATAG), and hPSMB9 (Forward: GCTTCACCACAGACGCTATT; Reverse: GCAGTTCATTGCCCAAGATG) were purchased from IDT.

### Cell Culture

Mammalian cells were all grown in a humidified incubator with 5% CO2 at 37°C. HEK293, Hs578T, MDA-MB-231, SW1573, A549 cells were cultured in Dulbecco’s Modified Eagle Media (DMEM, Gibco) supplemented with 10% FBS, 1% L-glutamine, and 1% penicillin and streptomycin. MM.1S, ANBL6, U266B1, RPMI-8226, 5TGM1-GFP, AHH1, MPC11, H460, H522, H1581 cells were grown in RPMI 1640 (Gibco) supplemented with 10% FBS, and 1% penicillin and streptomycin. MCF10A and 184B5 cells were cultured in DMEM/F-12 medium supplemented with 5% horse serum, 20 ng/ml EGF, 0.5 μg/ml hydrocortisone, 100 ng/ml cholera toxin, 10 μg/ml insulin and 1% penicillin and streptomycin. Transient transfection of HEK293 with mammalian expression vectors were carried out using FuGENE transfection reagent according to the manufacturer’s protocol. Cell lines were purchased from ATCC except 5TGM1-GFP (kind gift from Dr. Babatunde Oyajobi, University of Texas San Antonio, USA), MCF10A (kind gift from Dr. Alexandra Newton, University of California San Diego, USA) and ANBL6 (kind gift from Dr. Robert Orlowski, MD Anderson Centre, USA).

### Cytotoxicity assays

5,000-11,000 cells were plated per well in a 96-well plate. 4-6 h post plating, TIR-199 was added to the cells in biological triplicates at various concentrations ranging between 0-500 nM final concentration with 0 nM being DMSO-treated control. Cell viability was then determined using CellTiter 96^®^ AQueous Non-Radioactive Cell Proliferation Assay kit after 48 hours of drug treatment. Absorbance was measured in a Tecan multi-well plate reader and data represented as relative to DMSO control.

### CD138^+^ myeloma cell purification

CD138^+^ myeloma cells were purified from the bone marrow aspirates which were obtained from HIPAA compliant de-identified consenting patients in accordance with Institutional Review Board approved protocols at University of California-San Diego (UCSD). The bone marrow aspirates were kindly provided by Dr. Caitlin Costello, Moores Cancer Center, UCSD. CD138^+^ primary myeloma cells were purified from fresh bone marrow aspirates of multiple myeloma patients using EasySep™ Human Whole Blood and Bone Marrow CD138 Positive Selection Kit II (StemCell Technologies) following manufacturer’s instructions. Viable cells were collected for further analyses.

### Proteasome and immunoproteasome assays

For 26S constitutive proteasome activity assays, cells were lysed in 50 mM Tris/HCl (pH 7.5), 0.1% Nonidet P-40, 1 mM ATP, 10 mM MgCl2, 1:1000 β-mercaptoethanol and a phosphatase inhibitor cocktail (10 mM NaF, 20mM β-glycerophosphate). Proteasome peptidase activities from the cell lysates were assayed using fluorogenic peptide substrate Suc-LLVY-AMC (Enzo Life Sciences). The measured activity was normalized against total protein concentration. Fluorescence signal in whole cells or cell lysates were quantified using a Tecan Infinite^®^ M200 Pro multi-well plate reader.

Immunoproteasome activity assays were carried out as stated previously(32). Briefly, HEK293 cells transiently overexpressing FLAG tagged PSMB9 or PSMB8 were treated with IFNγ (100 ng/mL; 48 h) to induce expression of immunoproteasome. Cells were lysed in 50 mM Tris/HCl (pH 7.5), 0.1% Nonidet P-40, 1 mM ATP, 10 mM MgCl2, 1:1000 β-mercaptoethanol and a phosphatase inhibitor cocktail (10 mM NaF, 20mM β-glycerophosphate). Immunoproteasomes were affinity purified using FLAG agarose. Peptidase activities of purified immunoproteasomes were assayed using fluorogenic peptide substrates Ac-ANW-AMC for PSMB8 and Ac-PAL-AMC for PSMB9 (Boston Biochem). The measured activity was normalized against total protein concentration. Fluorescence signal in whole cells or cell lysates were quantified using a Tecan Infinite^®^ M200 Pro multi-well plate reader.

### Animal study

NSG mice (NOD.Cg-Prkdcscid Il2rgtm1Wjl/SzJ; Stock: 005557) were purchased from the Jackson Laboratory. Mice were housed and maintained at the University of California-San Diego (UCSD) in full compliance with policies of the Institutional Animal Core and Use Committee (IACUC). 5 × 10^6^ MM.1S cells resuspended in sterile PBS were injected into the tail-vein of 8-12 weeks old n=6 NSG mice of either sex. After 7 days, the mice were randomized into two groups n=3 each. TIR-199 was dissolved in 100% DMSO initially and further diluted down with 10 mM citrate buffer (pH 3.0) containing 10% (w/v) sulphobutylether-β-cyclodextrin (Carbosynth, USA) to a final DMSO concentration of 1%. Mice were injected at a daily intraperitoneal dose of 5 mg/Kg or vehicle alone for 4 consecutive days. 7 days after dose completion, mice were euthanized and both femurs were extracted rapidly, and fixed in 10% formalin.

### Micro Computed Tomography (μCT) imaging and data analysis

μCT imaging and data analysis was performed as stated previously(20) by investigators who were blinded to allocation. Femurs from MM.1S/NSG xenograft mice were fixed in 10% formalin for 24 hr and washed 2x in PBS. The femurs were imaged with a micro-computed tomography scanner, Skyscan 1076 (Kontich, Belgium). Each sample was wrapped in paper tissue that was moistened with PBS, and scanned at 9 μm voxel size, applying an electrical potential of 50 kVp and current of 200 μA, using a 0.5 mm aluminum filter. A beam hardening correction algorithm was applied prior to image reconstruction. Image data was visualized for each sample with Dataviewer and CTAn (Kontich, Belgium). A series of 2-D transaxial slices were generated over various regions of the femur. Fifteen slices were distributed around the center slice position with a separation of 0.06 mm for the metaphysis and mid-diaphysis regions, and 0.03 mm for the proximal femur region. Trabecular bone analysis was performed as appropriate for skeletally-mature animals(33). The trabecular region was selected by automatic contouring as close as possible to the periosteum but without overlapping the cortical bone. An adaptive threshold (using the mean maximum and minimum pixel intensity values of the surrounding ten pixels) was used to identify trabecular bone. From this region of trabecular bone, the tissue volume (24), trabecular bone volume (BV), trabecular bone volume fraction (% BV/TV), trabecular separation (Tb.Sp) and trabecular number (Tb.N) were determined. The technical quality of the image cross-sections was checked for damage before quantitative analysis was performed. Damaged areas were strictly avoided and never included in any quantitative analyses. Outliers were verified for legitimacy by checking the scan and reconstruction log file, image rotation, selection of the tip of growth plate, number of slices in metaphysis and diaphysis, and the contours.

### TIR-199 specificity profiling

TIR-199 specificity profiling assays were carried out at The International Centre for Protein Kinase Profiling (http://www.kinase-screen.mrc.ac.uk/). TIR-199 kinase specificity was determined against a panel of 50 protein kinases as described previously(34,35). The assay mixes and ^33^P-γATP were added by Multidrop 384 (Thermo). Results are presented as a percentage of kinase activity in DMSO control reactions. Protein kinases were assayed in vitro with 1 μM final concentration of TIR-199 and the results are presented as an average of triplicate reactions ± SD or in the form of comparative histograms.

### *In silico* docking studies

Molecular docking simulation was performed using AutoDock Vina software(36). AutoDock Tools (ADT) version 1.5.6 was used for structure pre-processing while Discovery Studio Visualizer and PyMOL were used to analyse bound conformations and interactions as reported previously(37). 3D structure of TIR-199 was drawn using ChemBio3D Ultra 11. To minimise energy, RMS gradient was set to 0.100 in each iteration. All structures were saved as SYBYL2 (.mol2) file format for input to ADT. Crystal structure of human immunoproteasome with all 14 subunits (PDB ID: 7AWE) was imported (as.pdb file) from the RCSB Protein Data Bank (PDB) (http://www.rcsb.org/). With the help of ADT, we extracted the crystal structure of PSMB8 and PSMB9 separately, bound ligand and water molecules were removed, and the polar hydrogens were added. Finally, Gasteiger charges were added to each atom and the non-polar hydrogen atoms were merged to the protein structure. The structure was saved in PDBQT file format.

### Statistics and data presentation

Details of all statistical tests and multiple comparisons used to derive P value has been detailed in Figure Legends. All experiments were repeated 2-3 times with multiple technical replicates to be eligible for the indicated statistical analyses, and representative image has been shown. All results are presented as mean ± SD unless otherwise mentioned. For animal studies, tumour-bearing mice of either sex were randomized into two equal groups of n=3 prior to vehicle or TIR-199 treatment. The investigators were blinded to allocation during μCT analysis and outcome assessment. Data were analysed using Graphpad Prism statistical package.

## Results

### TIR-199 inhibits the constitutive 26S Proteasome activity

TIR-199 (**Fig 1A**) is a syrbactin analog which has previously been reported as a proteasome inhibitor(30). TIR-199 potently inhibited the chymotryptic-like activity of immunoprecipitated PSMB5-FLAG with an IC50 is 50 nM (**Fig 1B**). To determine the efficacy of TIR-199-mediated inhibition of the endogenous proteasome, we treated HEK293 cells with 250 nM of TIR-199 for 6 hours and conducted immunoblots to assess if proteasome substrates would accumulate over time. Indeed, proteasome substrates p62, IKBα, and p21 protein levels increased suggesting that the proteasome has being inhibited by TIR-199 (**Fig 1C**). Furthermore, consistent with previous studies(30) we observed inhibition of chymotryptic-like endogenous proteasome activity in TIR-199-treated myeloma MM.1S cell lysates (**Fig 1D**). To further investigate the ability of TIR-199 to inhibit proteasome activity in primary refractory-patient-derived myeloma cells, we obtained bone marrow samples from 2 refractory multiple myeloma patients with extensive treatment history and recorded bortezomib-resistant relapse. We purified the CD138^+^ myeloma cells from the bone marrow treated these cells with either DMSO-control or TIR-199 at 500nM for 0.5 hours, and then measured the chymotryptic-like proteasome activity. Upon treatment with TIR-199, the proteasome activity decreased significantly compared to DMSO (**Fig 1D**), suggesting that TIR-199 is effective at decreasing proteasome activity in myeloma patients with extensive treatment history and potential bortezomib resistance. Furthermore, TIR-199 is specific and does not inhibit 50 other kinases tested in vitro even at 20-fold higher concentration to PSMB5 IC50 (**Supplementary Fig S1**).

**Figure 1.**
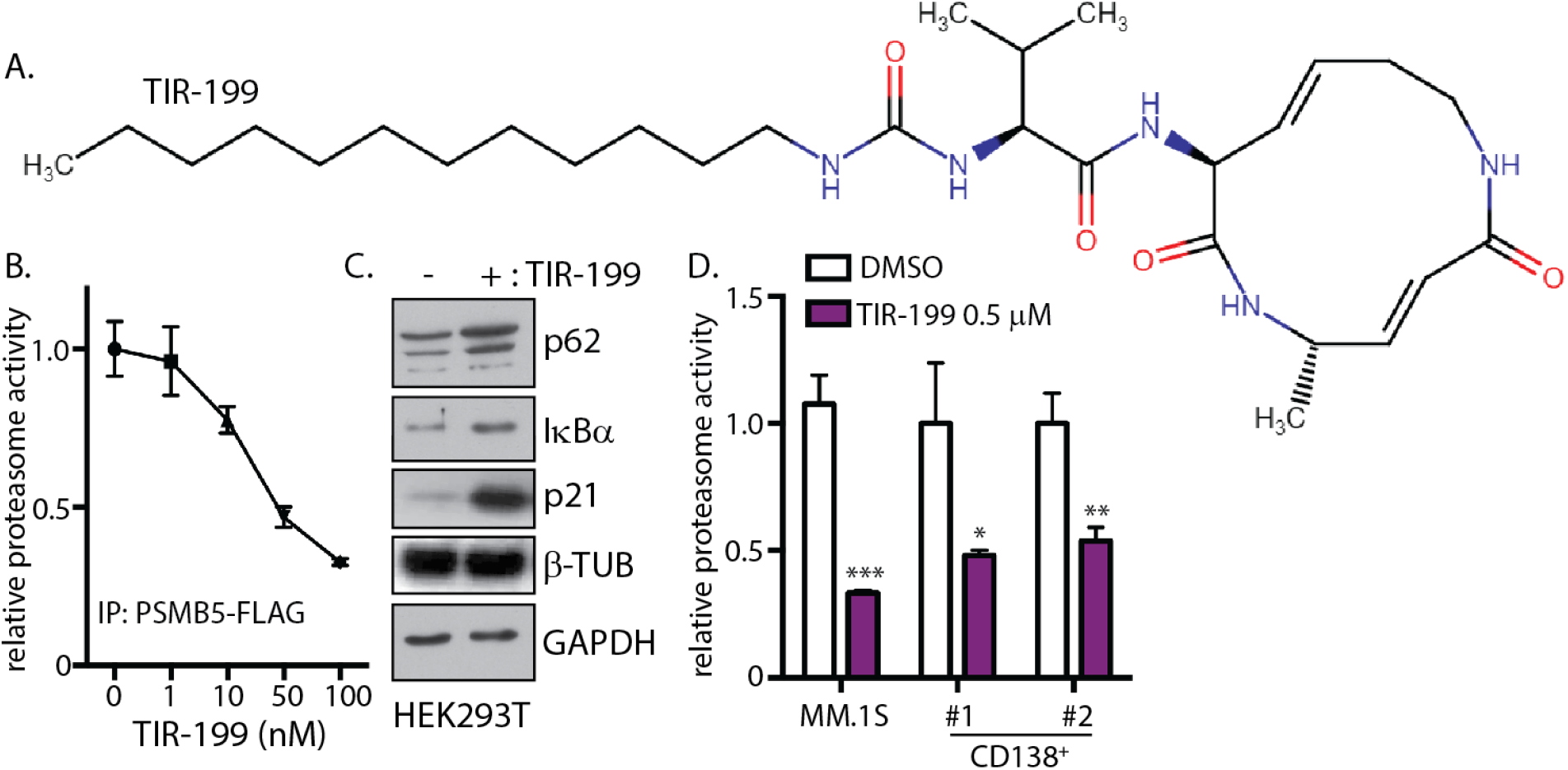
TIR-199 is a syrbactin based constitutive proteasome inhibitor. **(A)** Chemical structure of syrbactin derivative TIR-199. **(B)** Wild-type (WT) PSMB5-FLAG was immunoprecipitated from HEK293 cells stably expressing PSMB5-FLAG. PSMB5-FLAG was assayed using 100 μM Suc-LLVY-AMC in proteasome assay buffer with the indicated concentrations of TIR-199. Fluorescence signal upon release of AMC was measured in a Tecan plate reader. The results are presented as proteasome activity relative to the DMSO-treated control. Results are means ± S.D. for n=3 replicates. **(C)**HEK-293T cells were treated in the absence (DMSO) or presence of 0.5 μM of TIR-199 for 6 hr. Cell lysates were subjected to immunoblotting with the indicated antibodies. Similar results were obtained in two separate experiments. **(D)** Proteasome activity in lysates from MM.1S and primary CD138^+^ myeloma cells from two patients treated with DMSO or 0.5 μM TIR-199 for 0.5 h was measured with Suc-LLVY-AMC. *P < 0.05; **P < 0.01; ***P < 0.001 (compared to control treated, 2-tailed paired Student’s t test, mean ± SD from n = 3 replicates). Also refer to Supplementary Figure S1.

### TIR-199 Inhibits the Immunoproteasome

HEK293 cells ectopically expressing PSMB8 or PSMB9 were treated with IFNγ for 48 hours. A marked upregulation of PSMB8 and PSMB9 mRNA were observed which suggested immunoproteasome induction (**Fig 2A**). Flag agarose were utilised to immunoprecipitate PSMB8 or PSMB9 to pull-down the immunoproteasome. To determine how TIR-199 affects the immunoproteasome, activity assay was conducted on PSMB8 using Ac-ANW-AMC as the substrate peptide. Specific PSMB8 inhibitor ONX-0914(12) was used as a positive control for PSMB8 inhibition. Indeed, 100 nM TIR-199 addition inhibited PSMB8 potently and similarly to ONX-0914. However, like ONX-0914, TIR-199 at 100 nM did not inhibit PSMB9 activity suggesting a PSMB8-specific effect for TIR-199 (**Fig 2B**). To further predict TIR-199-binding to the immunoproteasome, *in silico* binding studies were conducted (**Fig 2C**). Docking analysis showed that TIR-199 formed six hydrogen bonds with Gly47, Thr1, Ser130, Glu116, Asp115 of PSMB8 with Thr1 forming two hydrogen bonds with the same bond length of 2.3 Å (**Supplementary Fig S2A**). Furthermore, 10 alkyl interactions through the dodecyl group of TIR-199 and multiple Van der Waals interactions enhanced the stability of the predicted TIR-199 bound PSMB8 structure (**Supplementary Fig S2A**). Interestingly, the predicted structure of TIR-199 and PSMB9 exhibited only 3 hydrogen bonds (**Supplementary Fig S2B**).

**Figure 2.**
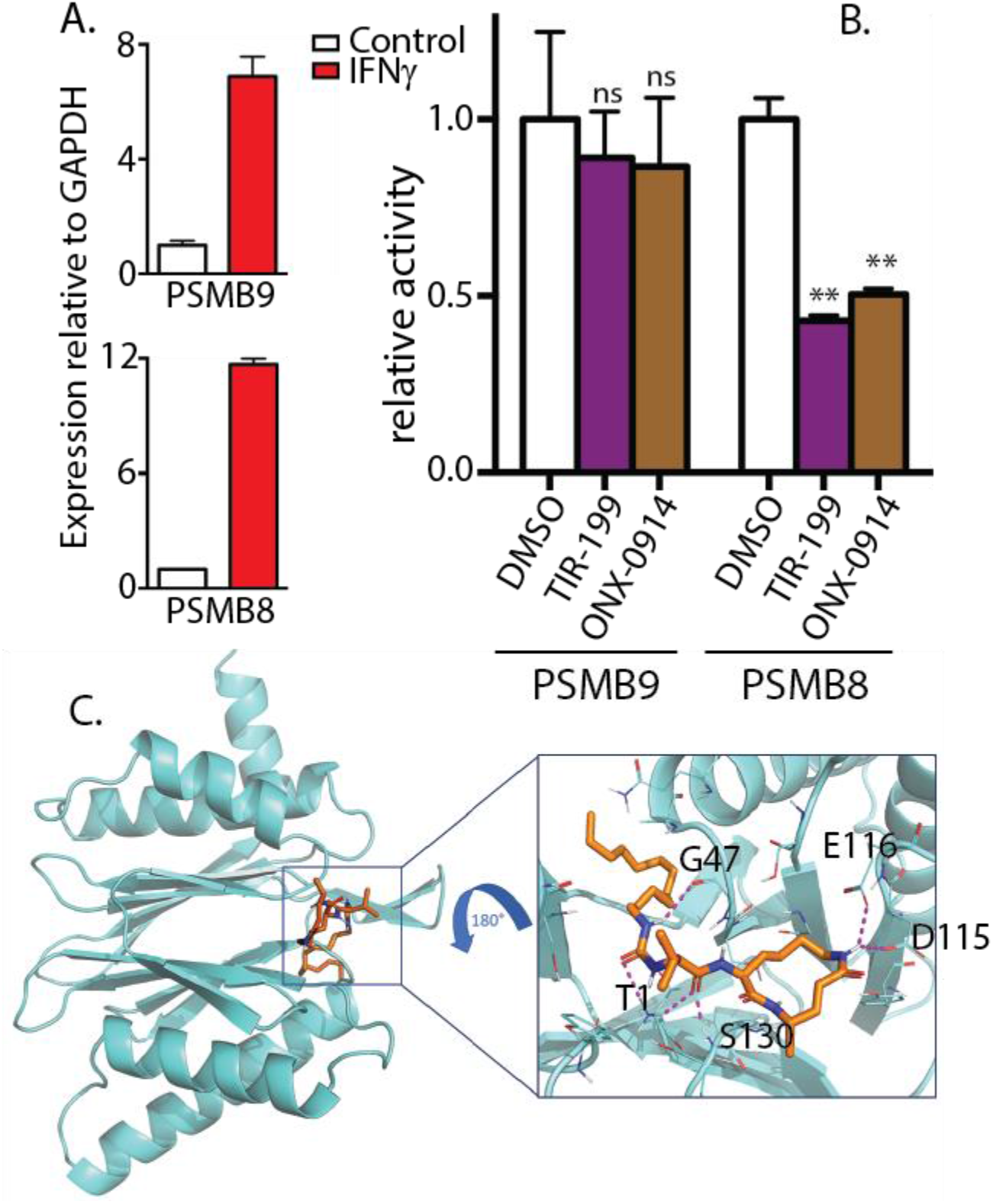
TIR-199 inhibits immunoproteasome subunit PSMB8 but not PSMB9. **(A)** Relative expression of PSMB8 and PSMB9 in HEK293 cells upon 100 ng/mL IFNγ stimulation for 48 hr. **(B)** PSMB8 or PSMB9 were immunoprecipitated from 48 hr 100 ng/mL IFNγ stimulated HEK293 cells transiently overexpressing either PSMB8-FLAG or PSMB9-FLAG. The FLAG immunoprecipitates were treated with either DMSO or 100 nM TIR-199 or 100 nM ONX-0914 for 30 min and activity assay were carried out with fluorogenic peptide substrates either Ac-PAL-AMC for PSMB9 or Ac-ANW-AMC for PSMB8. **P<0.01; ns: not significant (compared to DMSO control treated, 2-way ANOVA with Tukey’s multiple comparisons, mean ± SD from n=3 independent replicates). **(C)** TIR-199 (orange/blue) was docked into the crystal structure of human PSMB8 (cyan). Identified amino acid interacting residues are highlighted in black. Hydrogen bonds between TIR-199 and the amino acids in the binding pocket are shown by red dashed lines. Also refer to Supplementary Figure S2.

### TIR-199 causes cytotoxicity in cancer cells

It has been shown previously that TIR-199 can cause cytotoxicity in multiple myeloma cell lines MM1.S and U266(31). We conducted cytotoxicity assays on a panel of human and murine multiple myeloma cell lines (**Fig 3A**). Similar to most proteasome inhibitors reported till date, TIR-199 killed myeloma cells with an EC_50_ ranging between 1-50 nM. Similar to myeloma, triple negative breast cancer cells have also been reported to be particularly sensitive to proteasome inhibition(22). Therefore, we expanded the study to include TNBC cells and report the EC_50_ values for TIR-199 to be between 50-300 nM (**Fig 3B**). Although lung cancer cell lines have not been reported to be sensitive to proteasome inhibition, a case study did report exceptional therapeutic benefits of bortezomib in a KRAS mutated lung adenocarcinoma patient(24). Although within the lung cancer panel, EC_50_ ranged from 100 to >1000 nM for TIR-199, H460 was found to be the most sensitive lung cancer cell line to TIR-199 inhibition at 100 nM EC_50_ (**Fig 3C**).

**Figure 3.**
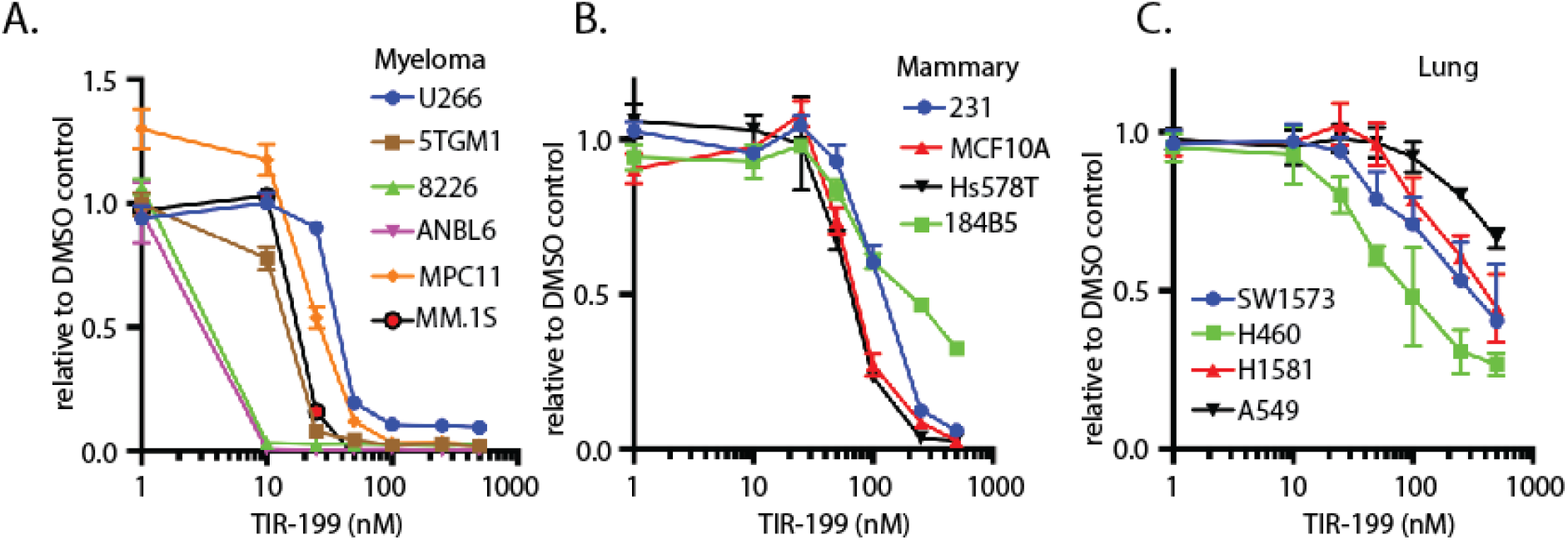
TIR-199 causes cell death. TIR-199 induces cytotoxicity in various **(A)** multiple myeloma, **(B)** triple-negative breast cancer, and **(C)** lung cancer cell lines after 48h incubation. MTS absorbance was measured in a Tecan plate reader. Cell viability was ascertained with CellTiter 96 AQueous Non-Radioactive Cell Proliferation Assay kit. Data were represented as percent viable compared to DMSO-treated cells. Results are means ± S.D. for n=3 replicates.

### TIR-199 bypasses PSMB5-mutation driven bortezomib-resistance and induces cytotoxicity in combination with DYRK2-inhibitor

Prolonged bortezomib treatment leads to development of drug-resistant disease relapse in multiple myeloma patients. Various mechanisms for bortezomib-resistance have been proposed and among them specific mutations in the PSMB5 subunit of the 20S proteasome core is thought to be one such mechanism. A recurrent bortezomib-resistant PSMB5 mutation observed in cells is the dual amino acid substitutions A49T and A50V (AA/TV), which directly affect bortezomib binding(38) and also reduces specific activity of the proteasome(8). Since TIR-199 have an altered binding mechanism to PSMB5 compared to bortezomib and does not exhibit direct hydrogen bond interaction with Ala49 (27), we hypothesized that AA/TV harbouring cells will exhibit similar sensitivity to TIR-199 as parental cells. We generated HEK293 cells with stable ectopic overexpression of wild-type (WT) or AA/TV PSMB5 with a C-terminal FLAG tag. We utilized FLAG tag pull down to immunoprecipitate the 26S proteasome and observed similar proteasomal subunit distribution on a Coomassie-stained gel for both WT and AA/TV (**Fig 4A**). When we further assayed proteasome activity on the immunoprecipitates using the peptide substrate Suc-LLVY-AMC, we observed a reduced proteasome activity in AA/TV mutant compared to WT (**Fig 4B**) which is consistent with previous report. In order to determine sensitivity of AA/TV PSMB5 toward TIR-199, the immunoprecipitated PSMB5 was treated with either TIR-199 or bortezomib. Treatment of WT PSMB5 with 10 nM bortezomib led to a dramatic reduction in proteasomal activity while AA/TV mutant exhibited significant resistance (**Fig 4C**). However, both WT and AA/TV harbouring cells were equally sensitive to 100 nM TIR-199 (**Fig 4C**). This suggests that TIR-199 binds to the constitutive proteasome via a different mechanism than bortezomib, allowing it to bypass bortezomib resistance. Interestingly, a combination of TIR-199 with DYRK2-inhibitor LDN192960 exhibited a modest additive effect toward the biochemical inhibition of proteasome activity in MM.1S cells (**Fig 4D**). Next, we wanted to determine if LDN192960-TIR-199 mediated additive impairment of proteasome activity could have an effect on cancer cell viability. Interestingly, a remarkable synergistic cytotoxicity was induced in proteasome addicted MM cell lines MPC11 and 5TGM1 upon treatment with LDN192960 and TIR-199 (**Fig 4E**).

**Figure 4.**
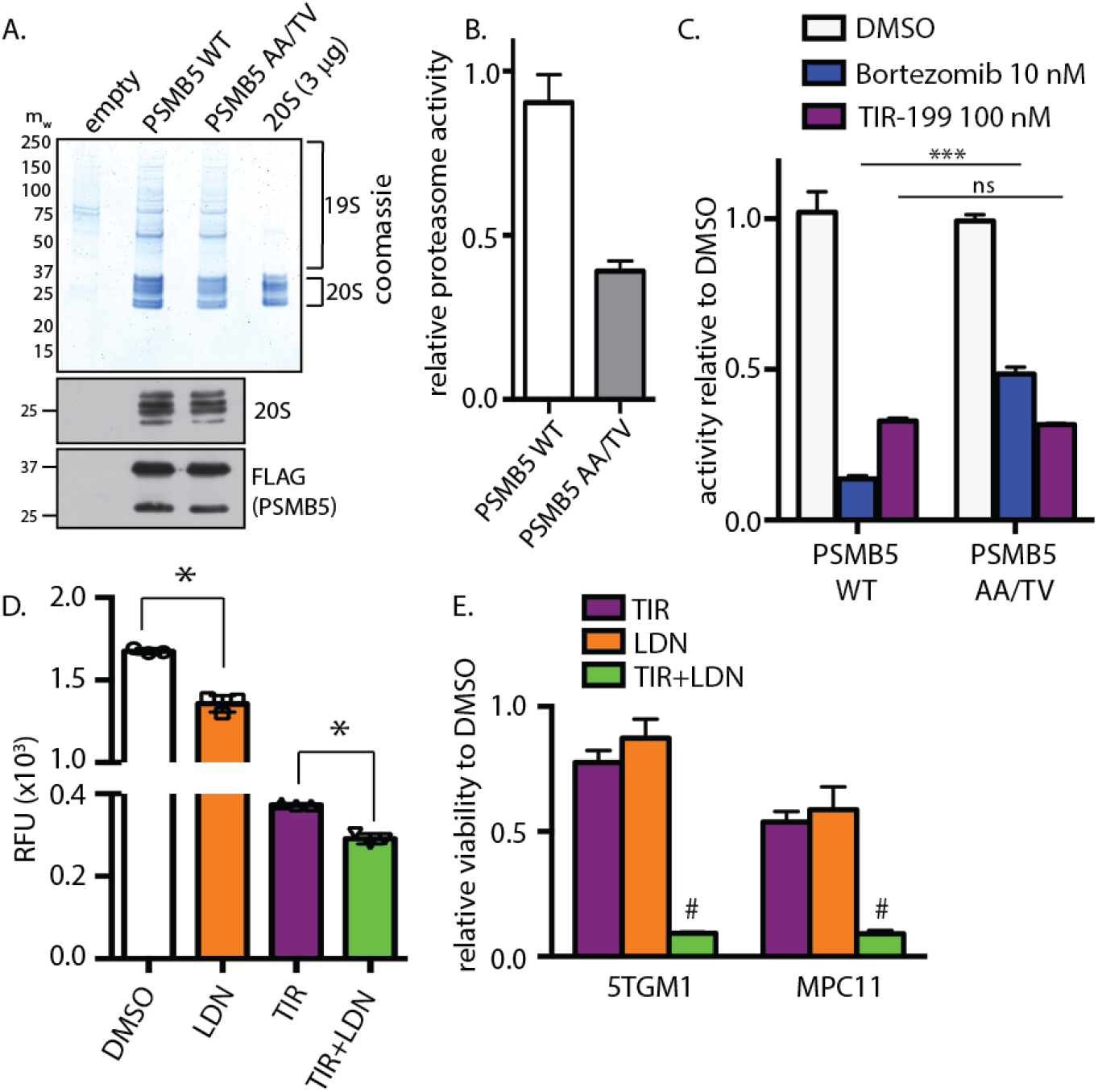
TIR-199 bypasses bortezomib-resistant PSMB5 AA/TV mutation and induces synergistic cytotoxicity in myeloma cells with LDN192960. **(A)** WT and AA/TV mutated PSMB5-FLAG were immunoprecipitated from HEK-293 cells stabling expressing the wild-type and mutant enzymes and were analysed by Coomassie Blue staining of a polyacrylamide gel (top panel) and immunoblotting with indicated antibodies (bottom panel). Purified 20S proteasome was loaded as control to predict amount. **(B)** Intrinsic proteasome activities of the equivalent amounts of PSMB5 and PSMB5(AA/TV) were compared by carrying out a proteasome activity assay using Suc-LLVY-AMC substrate peptide. Values are means±S.D. for an experiment carried out in triplicate. **(C)** Proteasome activities of the equivalent amounts of PSMB5 and PSMB5(AA/TV) were compared in the presence or absence of indicated molecules by carrying out a proteasome activity assay using Suc-LLVY-AMC substrate peptide. Data represented as activity relative to respective DMSO-treated sample. ***P<0.001; ns: not significant (compared to DMSO control treated, 2-way ANOVA with Tukey’s multiple comparisons, mean ± SD from n=3 independent replicates). **(D)** MM.1S cells were pretreated with indicated drugs or combination (TIR, TIR-199; LDN, LDN192960) for 1 h, and proteasome activity was measured in cell lysates using Suc-LLVY-AMC. *P < 0.05 (compared to control treated, ordinary 1-way ANOVA, mean ± SD from n = 3 independent experiments). **(E)** Multiple myeloma cells MPC11 and 5TGM1 were treated with TIR-199 alone (10 nM for 5TGM1 and 25 nM for MPC11) or with LDN192960 alone (3 μM for 5TGM1 and 5 μM for MPC11) or the combination of TIR-199 and LDN192960 for 48 h, and cell viability was analyzed by CellTiter 96 AQueous Non-Radioactive Cell Proliferation Assay kit. Viability of DMSO-treated cells was utilized as control. Data are represented as fold viability of DMSO-treated control for each cell line (# indicates statistical significance compared to each single drug treatment; 2-way ANOVA with Sidak’s multiple comparison)

### TIR-199 impedes multiple myeloma progression in vivo

Previous studies have shown that TIR-199 is active in vivo and can reduce cancer cell proliferation in hollow fiber assays(30) and an ectopic tumour xenograft model(31). Although TIR-199 can reduce ectopic tumours, no information is available regarding the ability of TIR-199 in targeting systemic myeloma-mediated bone degeneration in vivo which is the major hallmark of multiple myeloma. To address this, we intravenously injected MM.1S cells into NSG mice of either sex to generate a systemic multiple myeloma model exhibiting the canonical myeloma-induced systemic bone degeneration. Mice were randomized one week post intravenous MM.1S implantation and n=3 mice each were treated with either vehicle control or TIR-199 at 5mg/Kg per day for 4 consecutive days. We used a lower dose keeping in mind the relatively toxic effects of systemic proteasome inhibition. 7 days post treatment-completion the femurs were collected, fixed and μCT quantitative analysis was performed on the femur pairs of 3 vehicle-treated and 3 TIR-199-treated mice (**Fig 5A**). Trabecular bone was quantified in the distal metaphysis and proximal regions independently and data was presented by combining both analysis for each cohort to provide a more holistic view. Quantitative analysis of the trabecular bone in both the distal and proximal regions of the femurs presented a higher trabecular number (**Fig 5B**) which is the number of traversals across a trabecular region per unit length on a random linear path through the volume of interest. The TIR-treated mouse femurs also exhibited significantly lower trabecular separation (**Fig 5C**) which is the thickness of spaces between trabecular bone. Although the combined distal and proximal numbers for % bone volume over total volume was not significant (**Fig 5D**), the TIR-199-treated mouse femurs had less damage. This suggests that TIR-199 treatment at a low dose of 5mg/Kg for 4 days has slowed down the myeloma-mediated bone degeneration process.

**Figure 5.**
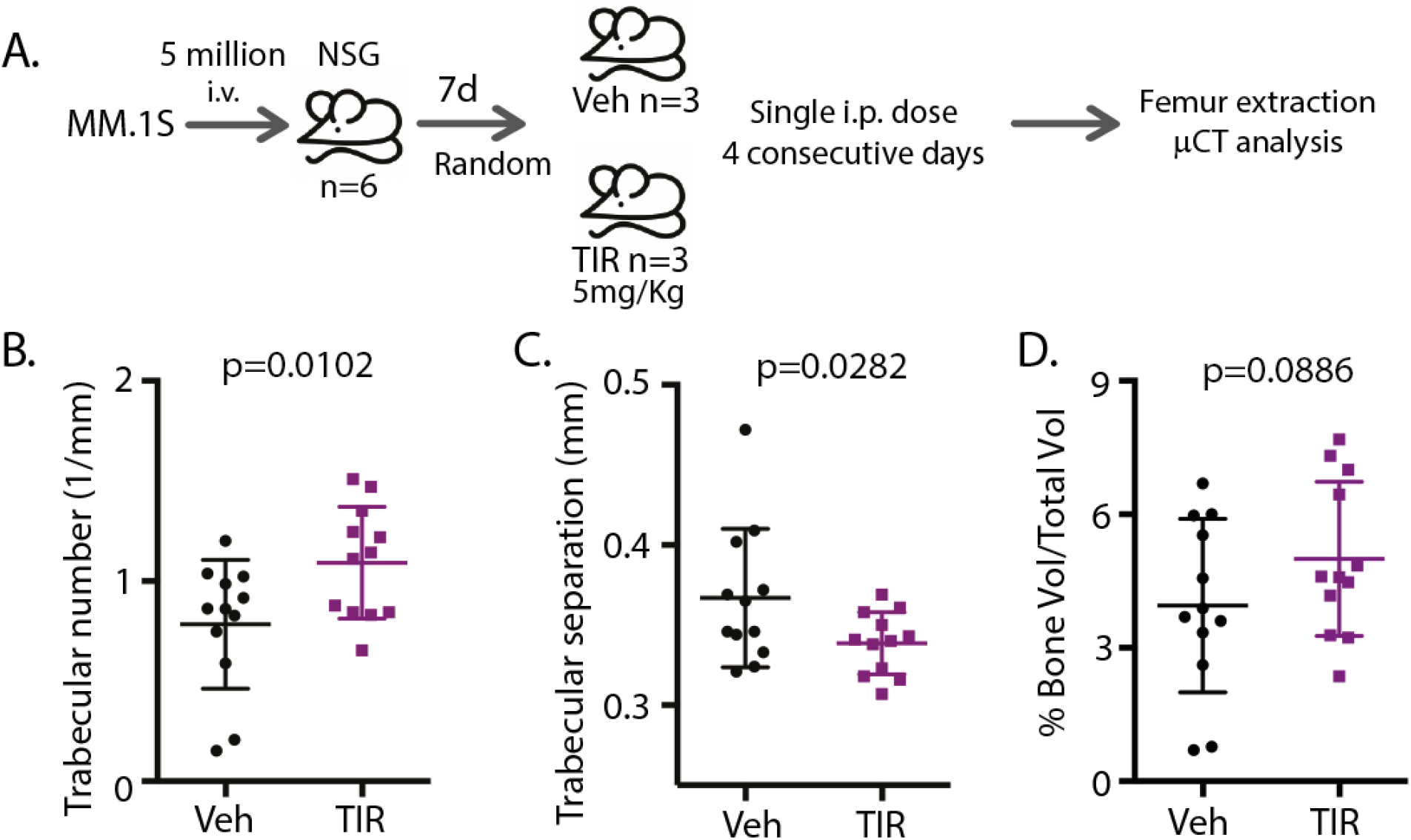
Low dose TIR-199 delays myeloma-mediated bone degeneration. **(A)** MM.1S cells were injected i.v. into NSG mice n = 6. One-week postinjection, mice were randomized into 2 groups of n = 3 and treated with vehicle or TIR-199 5 mg/kg i.p. daily for 4 consecutive days. One-week posttreatment, the mice were killed, and formalin-fixed femur bones were imaged using μCT. Post-μCT the trabecular number **(B)**, the trabecular separation **(C),** and percentage of bone volume over total volume **(D)** were quantified for distal and proximal femurs. P-value shown for vehicle versus TIR-199-treatment and derived using t-test. Data represented as mean ± SD, from n = 6 femurs combining distal and proximal regions’ analyses for each.

## Discussion

In the current study we extensively characterize syrbactin-class molecule TIR-199 as a potent constitutive proteasome **(Fig 1)** and PSMB8 immunoproteasome subunit inhibitor **(Fig 2)**. We further establish that TIR-199 treatment indeed leads to accumulation of proteasome substrates in cells **(Fig 1C)** and potently reduce proteasome activity in relapsed and refractory patient-derived primary myeloma cells **(Fig 1D)**. Together with the ability to bypass A49T+A50V PSMB5 mutations **(Fig 4C)** suggests that TIR-199 can bypass bortezomib drug-resistance which is a recurrent and global problem in the field of myeloma therapeutics. Unlike reversible boronates or irreversible epoxyketones, previous co-crystallography studies showed that syrbactin based molecules use a covalent interaction with PSMB5 Thr1 residue to stabilize the inhibitor bound structure(27). Our *in silico* docking studies with immunoproteasome subunits predicted that Thr1 on PSMB8 is also involved in binding to TIR-199 **(Fig 2C**). However, bortezomib binding involves a critical hydrogen bond with Ala49 on PSMB5 (38) which is not observed in case of TIR-199 **(Fig 2C and supplementary Fig S2A)** and this explains the mechanism by which TIR-199 is able to bypass bortezomib resistance in the AA/TV PSMB5 mutation **(Fig 4C).** Indeed, a previous study has reported bortezomib-resistant myeloma and mantle-cell lymphoma cells were sensitive to TIR-199(31). Furthermore, TIR-199 clearly prefers the PSMB8 subunit over PSMB9 (**Fig 2B**) and this is probably due to the reduced number of hydrogen bonds predicted for TIR-199-PSMB9 interaction (**Supplementary Fig S2B**). Inhibition of dual classes of proteasomes by TIR-199 could bode well for patients since epoxyketone-class dual inhibitor oprozomib is currently showing promise in clinical trials(11). TIR-199 could expand the repertoire of proteasome-based therapeutics since further drug resistance is expected to appear for the next generation of boronates and epoxyketones as well. The ability of TIR-199 to induce cytotoxicity in multiple myeloma, TNBC and lung cancer lines (**Fig 3**) further attests to its potency and the fact that the molecule can synergistically induced myeloma cell death in combination with DYRK2 inhibitor LDN192960 (**Fig 4D&E**) further opens new possibilities in therapeutic targetting of proteasome regulators. Proteasome addiction is well documented for myeloma and TNBC cells, while more recently, a small proof-of-concept human trial showed that some lung adenocarcinoma cancer patients with KRAS G12D mutation responded well to proteasome inhibitor therapy(24). In our study, we found that lung cancer cell line H460, which harbors a KRAS Q61H mutation, was maximally sensitive to TIR-199 within the lung cancer panel (**Fig 3C**). Paradoxically, KRAS G12C harboring A549 cells seemed relatively more resistant (**Fig 3C**). This suggests that proteasome targetting may have a broader therapeutic potential than previously appreciated and further work is needed to establish a potential link between KRAS mutations and proteasome-inhibitor sensitivity.

Various kinases have been reported to regulate proteasome activity including DYRK2(14), PIMs(17), PKA(15), and PKG(16). In fact PIM kinases are currently being pursued as active targets for multiple myeloma(39) and it would be ideal to study the cytotoxic potency of TIR-199 in combination with similar proteasome-regulating kinase inhibitors to ascertain the ideal therapeutic regimen for the future. As a monotherapy option, TIR-199 has been shown to reduce ectopic tumour volume previously(31) while we show that dosing even at one-fifth the maximal tolerated dose, TIR-199 can significantly delay systemic myeloma-mediated bone degeneration in an orthotopic mouse xenograft model (**Fig 5**). Mice treated with TIR-199 showed improved trabecular bone numbers (**Fig 5B**) and smaller gaps between the bones (**Fig 5C**) suggesting improved bone health compared to vehicle-treated control and consequently reduced myeloma burden. Since we were limited by drug availability, we could not study the beneficial effect of TIR-199 treatment on overall survival over a longer period, however, our data clearly suggests that TIR-199 exhibits all the bona fide hallmarks of a novel class of proteasome inhibitor.

With over 160,000 patients worldwide, multiple myeloma accounts for nearly 10% of all hematological malignancies (40). Patients exhibit extensive drug resistance with high therapy-induced toxicities(7,13). Hence, establishing novel syrbactin class TIR-199 could indeed expand the therapeutic option for refractory patients and extend their survival considerably. Further work is needed to translate syrbactin-class proteasome inhibitors to the clinic, however, the lead compound TIR-199 is indeed a promising drug for the near future.

## Data availability statement

Data and reagents are available upon request to the corresponding author SB. TIR-199 has been licensed to Zymergen Inc., Emeryville, California, USA and not available from the authors.

## Acknowledgement

This work was supported by the Cancer Research UK EDDPMA-May21\100005 (to SB), Ninewells Cancer Campaign Cancer Research Award (to SB), Royal Society RGS\R2\212056 (to SB) and University Grants Commission F-540/5/CASII/2018 SAP-I (to HMP). The authors thank Dr Wen Ma for valuable input and Joseph Maroge for coordinating with consenting patients and transferring samples. We thank the Kinase-Screen team at International Centre for Protein Kinase Profiling, Dundee, UK.

---

**Supplementary Figure S1.**
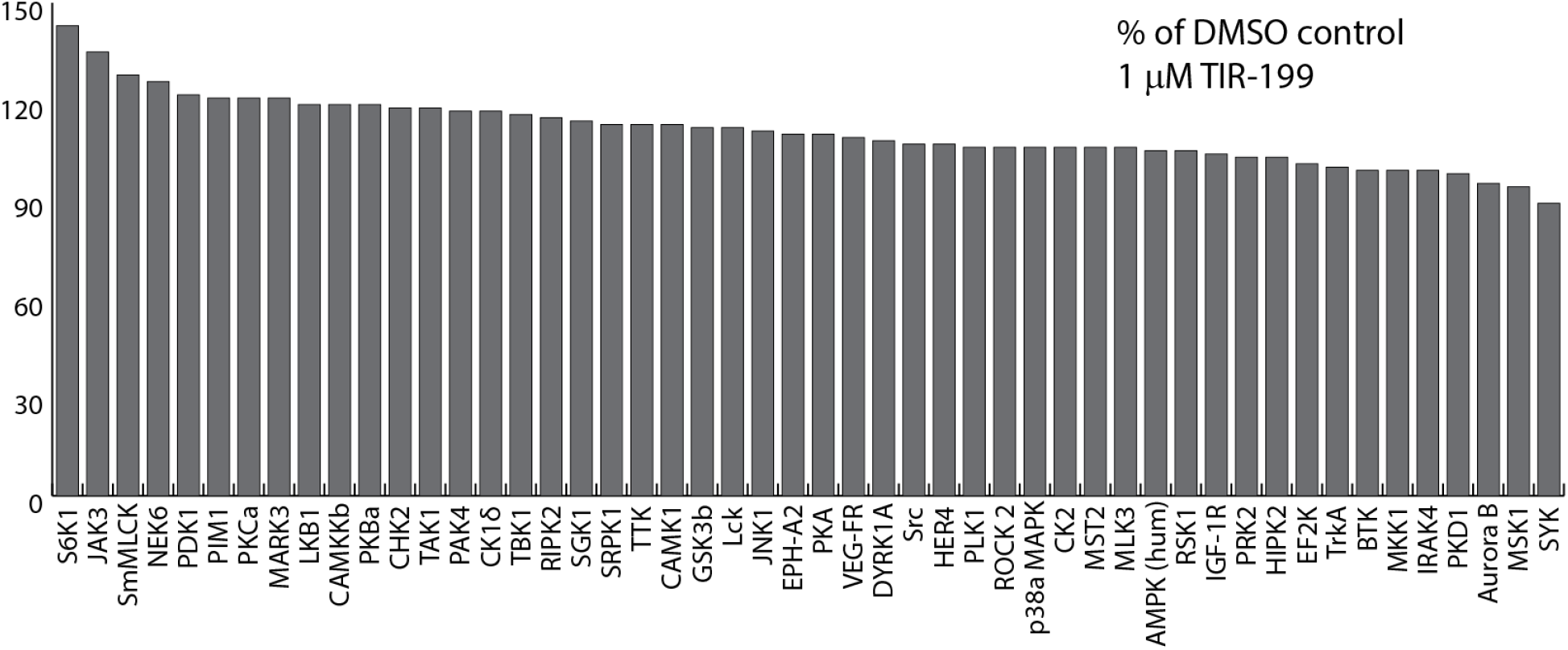
TIR-199 does not inhibit the activity of 50 kinases in vitro. Kinase profiling of TIR-199 at 1 μM was carried out against the panel of 50 kinases at the International Centre for Protein Kinase Profiling (http://www.kinase-screen.mrc.ac.uk/).

**Supplementary Figure S2.**
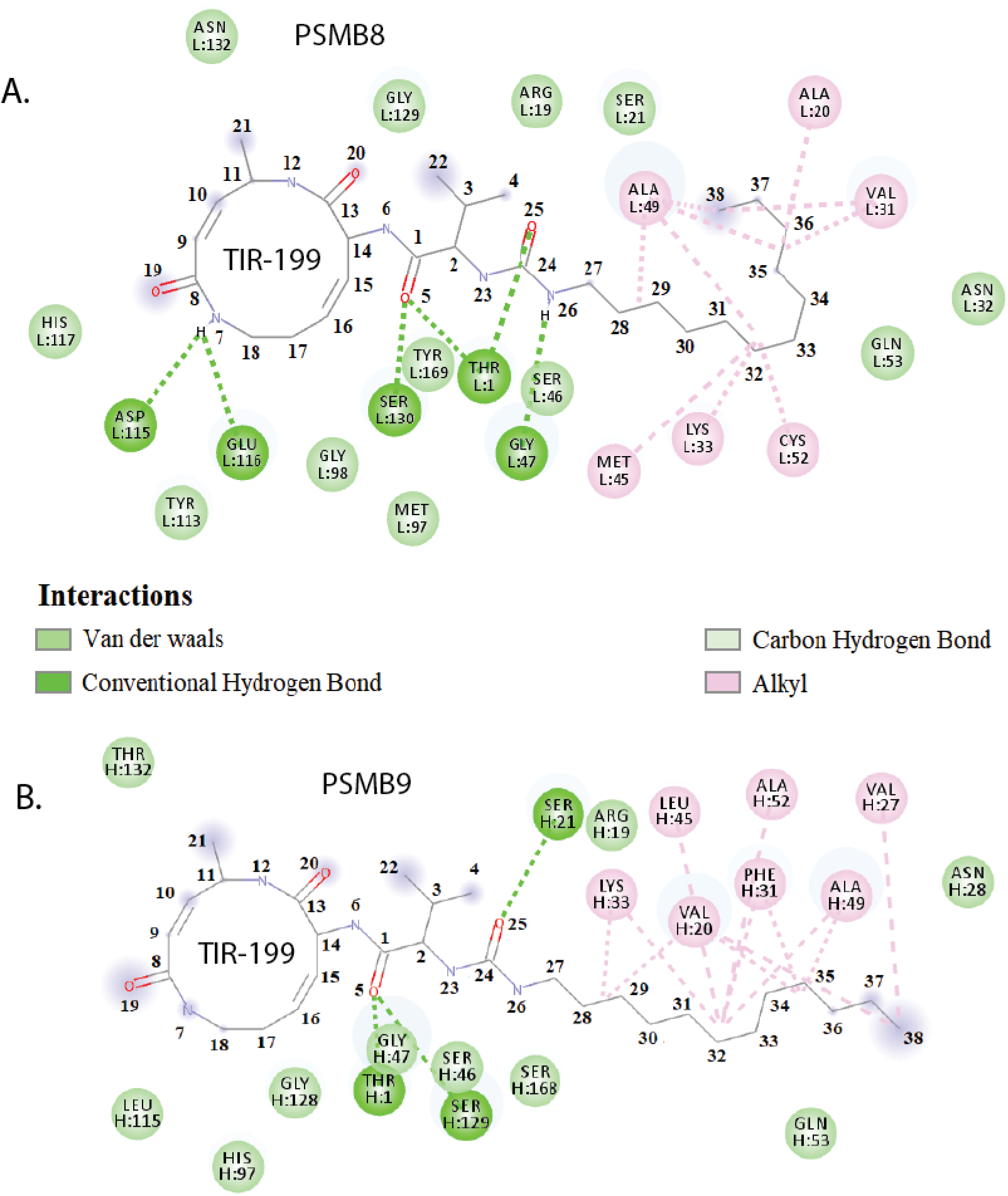
TIR-199 is predicted to form 3 more hydrogen bonds with PSMB8 over PSMB9. 2D interaction diagram showing TIR-199 docking mechanism in the active site of PSMB8 **(A)** and PSMB9 **(B).**

